# Widefield Acoustics Heuristic: Advancing Microphone Array Design for Accurate Spatial Tracking of Echolocating Bats

**DOI:** 10.1101/2025.06.03.657701

**Authors:** Ravi Umadi

**Affiliations:** Lehrstuhl für Zoologie, TUM School of Life Sciences, Technical University of Munich, Liesel-Beckmann Str 4, 85354, Freising, Germany

**Author notes:** Corresponding author: Ravi Umadi–.

**Keywords:** Microphone 3D Array, Bat Call Localisation, Echolocation, Array Design, TDoA

## Abstract

Accurate three-dimensional localisation of ultrasonic bat calls is essential for advancing behavioural and ecological research. I present a comprehensive, open-source simulation framework—*Array WAH* —for designing, evaluating, and optimising microphone arrays tailored to bioacoustic tracking. The tool incorporates biologically realistic signal generation, frequency-dependent propagation, and advanced Time Difference of Arrival (TDoA) localisation algorithms, enabling precise quantification of both positional and angular accuracy. The framework supports both frequency-modulated (FM) and constant-frequency (CF) call types, the latter characteristic of Hipposiderid and Rhinolophid bats, which are particularly prone to localisation errors due to their long-duration emissions. A key innovation is the integration of source motion modelling during call emission, which introduces Doppler-based time warping and phase shifts across microphones—an important and often overlooked source of error in source localisation.

I systematically compare four array geometries—a planar square, a pyramid, a tetrahedron, and an octahedron—across a volumetric spatial grid. The tetrahedral and octahedral configurations demonstrate superior localisation robustness, while planar arrays exhibit limited angular resolution. My simulations reveal that spatial resolution is fundamentally constrained by array geometry and the signal structure, with typical localisation error ranging between 5–10 cm at 0.5 m arm lengths.

By providing a flexible, extensible, and user-friendly simulation environment, *Array WAH* supports task-specific design and deployment of compact, field-deployable localisation systems. It is especially valuable for investigating the acoustic behaviour of free-flying bats under naturalistic conditions, and complements emerging low-power multichannel ultrasonic recorders for field deployment and method validation.

## 1 INTRODUCTION

Echolocating bats exhibit remarkable diversity in their biosonar strategies. Species of the Vespertilionid family produce short, broadband frequency-modulated (FM) calls that are highly effective for spatial localisation in cluttered environments. In contrast, members of the Hipposiderid and Rhinolophid families emit long-duration constant-frequency (CF) calls, enabling precise detection of fluttering prey and Doppler shift analysis. Regardless of call type, echolocation behaviour is inherently dynamic and context-dependent. Bats actively adjust call structure, repetition rate, and beam direction in response to changing sensory demands and ecological constraints [1–8]. This behavioural flexibility, while critical for efficient navigation and prey capture, complicates acquiring accurate behavioural and spatiotemporal data needed to understand bat ecology and neuroethology.

Laboratory studies, while providing controlled settings, often fail to replicate the full range of natural behaviours due to spatial restrictions and artificial environments [7]. Therefore, field-based acoustic localisation using microphone arrays is crucial. However, practical challenges arise: ultrasonic bat calls typically span frequencies between 20 and 100 kHz, with strong attenuation and directionality in air [9–12], making robust localisation technically demanding. Furthermore, the diversity among bat species implies a wide range of call parameters and behaviours, necessitating flexible and adaptable localisation systems.

Summary. Bats, belonging to the order Chiroptera, exhibit extraordinary diversity in both ecological specialisation and behavioural traits—especially in their use of echolocation. The variety in call structures, adaptive strategies, and sensory behaviours makes them one of the most intriguing subjects in bioacoustic research. Tracking their movements using echolocation calls offers powerful insights for both scientific study and conservation efforts. However, working with ultrasound presents significant technical challenges, often requiring bulky and costly multi-microphone arrays that demand specialised expertise. While efforts are underway to develop lightweight and portable systems, three-dimensional (3D) microphone arrays offer unique advantages in terms of spatial coverage and compactness.

To support this progress, I developed Array WAH—a software toolkit that enables researchers to evaluate and compare different 3D microphone array geometries. The tool simulates acoustic localisation scenarios and visualises spatial accuracy patterns, making it easier to assess array performance before physical development and deployment. By bridging the gap between conceptual design and field-ready hardware, Array WAH helps accelerate the development of effective and efficient acoustic tracking systems for use in bat research and beyond.

Existing microphone arrays in bioacoustics largely rely on planar two-dimensional configurations [13]. While easier to build, these 2D arrays are limited by spatial ambiguities and suffer from degraded localisation accuracy over large volumes. For precise localisation in 3D, the planar arrays need to span wider. Three-dimensional volumetric arrays can mitigate these limitations, offering enhanced spatial resolution and directional precision within smaller physical footprints. Nonetheless, their design and optimisation are hindered by the lack of accessible tools capable of simulating the complex acoustic propagation and localisation processes at ultrasonic frequencies.

In parallel, acoustic localisation methods such as time difference of arrival (TDoA) and steered response power (SRP) [14] algorithms have been employed successfully in various contexts [15, 16]. However, the efficacy of these methods depends strongly on array geometry, signal frequency content, environmental conditions, and algorithmic assumptions. This interplay highlights the critical need for a systematic framework to evaluate array performance before field deployment.

While Acoustic Vector Sensors (AVS) combined with Acoustic Intensity Measurement (AIM) methods have demonstrated effectiveness in localising bird calls [17] and other lower-frequency animal vocalizations [18, 19], their direct application to bat echolocation presents significant challenges. The ultrasonic braodband bat calls would require sensors with extremely fine spatial resolution and high sensitivity to accurately capture particle velocity and acoustic intensity, which conventional AVS designs are not optimized for. Moreover, ultrasonic calls suffer from rapid atmospheric attenuation and complex directional beam patterns, complicating the assumptions underlying AIM algorithms, which rely on relatively stable intensity fields and narrowband signals. As a result, AVS and AIM techniques, although powerful in other bioacoustic contexts, are generally unsuitable for precise three-dimensional localisation of bat calls, necessitating the development of specialised microphone array configurations and localisation algorithms tailored to the unique acoustic properties of bat biosonar.

Addressing this gap, I developed *Array WAH* (Widefield Acoustics Heuristic) [20], a versatile simulation and analysis platform tailored for ultrasonic bat call localisation. The term *Widefield* refers to the broad three-dimensional spatial coverage achieved by the volumetric microphone arrays, enabling comprehensive localisation of ultrasonic bioacoustic signals across extended volumes typical of behavioural field studies. *Array WAH* integrates detailed signal generation based on empirically inspired FM and CF bat calls, models frequency-dependent atmospheric attenuation, motion induced Doppler shifts, and simulates multi-microphone signal reception incorporating realistic propagation delays and amplitude scaling. The framework further applies TDoA-based multilateration for source localisation and supports spatial grid sweeps to map positional and angular errors across volumes representative of natural bat flight spaces.

In this study, I employed the *Array WAH* software toolkit to analyse four distinct microphone array configurations: three four-microphone geometries (tetrahedral, planar square, and pyramidal) and one six-microphone octahedral array. These standard geometries were chosen to simplify the description and allow clear comparison of localisation performance across different spatial layouts. The primary aim was to demonstrate the core principles and capabilities of the method rather than to optimise array design for a specific field application. Notably, the *Array WAH* toolkit allows users to define and evaluate arbitrary array geometries, making it a flexible resource for custom, task-specific localisation system development.

The results reveal that volumetric (3D) arrays, such as the tetrahedral and pyramidal layouts, consistently outperform planar arrangements in spatial precision. Specifically, volumetric arrays offer enhanced localisation accuracy within target volumes on the order of 5 m^3^, typical of experimental behavioural studies with bats. The *Array WAH* toolkit facilitates this assessment through the generation of detailed spatial error 3D plots, heatmaps and comprehensive statistical summaries, equipping researchers with actionable insights to tailor microphone arrays for portability and deployment ease without compromising precision.

Ultimately, *Array WAH* empowers bioacousticians to systematically design, simulate, and validate microphone arrays optimized for the accurate localisation of high-frequency ultrasonic signals. This capability addresses critical technical challenges in field-based studies of bat echolocation, thereby advancing research in behavioural ecology and conservation biology.

## 2 METHODS

### 2.1 Simulation Framework

I developed a MATLAB-based simulation framework to evaluate the accuracy of three-dimensional acoustic source localisation across different microphone array geometries. The core implementation is encapsulated in a custom object-oriented class, BatCallLocaliser, which integrates signal generation, propagation modelling, TDoA extraction, multilateration, and result logging. Figure 1 provides an overview of the framework workflow.

**Figure 1:**
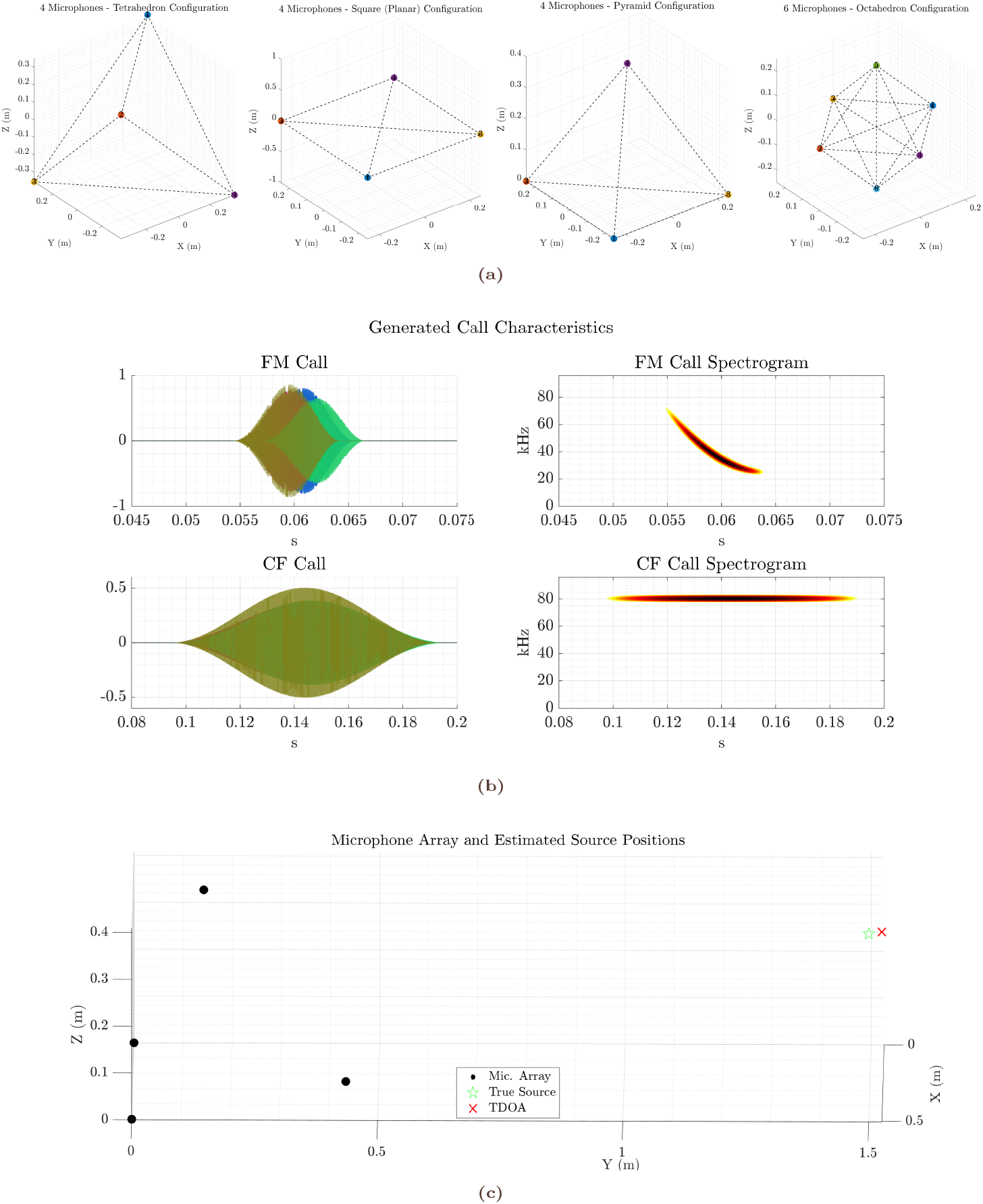
Methodological overview of microphone array configurations and bat call simulation. **(a)** 3D visualizations of the four microphone array geometries used in this study: Tetrahedron, Square (Planar), Pyramid (4microphones) and Octahedral (6-microphones) configurations. Each colored point represents a microphone, connected by dashed lines illustrating array topology. **(b)** Representative synthesised bat call waveforms (left; each colour indicates a microphone channel) and the corresponding spectrogram (right) from one of the channels. The top row illustrates a frequency-modulated (FM) sweep characteristic of vespertilionid echolocation signals, while the bottom row shows a constant-frequency (CF) call typical of Hipposiderid and Rhinolophid bats. These examples demonstrate the distinct spectral-temporal signatures of each call type as simulated across the 4-channel array. **(c)** Example spatial plot showing the relative positions of microphones (black dots), the true bat call source location (green star), and the source location estimated by time difference of arrival (TDoA) method (red cross).

#### Microphone Array Configurations

Four standard geometric microphone array configurations were tested:

1. **Tetrahedral array**: Four microphones placed at the vertices of a regular tetrahedron.
2. **Planar square array**: Four microphones forming the corners of a square in a single horizontal plane.
3. **Pyramidal array**: A square base with three microphones and a fourth at the apex, forming a square pyramid.
4. **Octahedral array**: Six microphones arranged at the vertices of a regular octahedron, offering higher spatial symmetry.

For each configuration, the array geometry was parameterised by an edge length variable *d*_edge_, defining the spatial scale. Microphone coordinates were specified in a *N ×* 3 matrix **M**, where each row corresponds to the 3D position of a microphone. These configurations are illustrated in Figure 1a.

#### Virtual Bat Call Generation

To simulate echolocation signals emitted by flying bats, I synthesised two types of source signals: frequency-modulated (FM) sweeps and constant-frequency (CF) tones. Both types optionally incorporate Doppler compression or stretching to mimic source motion effects due to the bat’s relative velocity.

##### FM Calls

The FM call was constructed as a quadratic downward chirp of duration *d*, starting at frequency *f*_0_ and ending at *f*_1_. The unwarped signal is defined as:

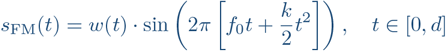

where 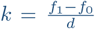 is the chirp rate, and *w*(*t*) is a Hanning window applied to reduce spectral leakage.

To simulate motion-induced frequency distortion, the waveform was time-warped according to the Doppler shift factor:

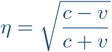

where *v* is the bat’s radial velocity (positive if receding), and *c* = 343 m*/*s is the speed of sound. Doppler distortion was applied via rational resampling using the factor *η*, thereby compressing or stretching the waveform duration depending on motion direction.

##### CF Calls

Constant-frequency tones were generated as:

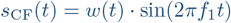

using a windowed sinusoid of frequency *f*_1_, duration *d*, and Hanning envelope *w*(*t*). Doppler resampling identical to the FM case was applied to simulate tone warping under source motion.

##### Post-processing

To better reflect natural emission patterns, each signal was zero-padded on both ends by a percentage defined by a tail parameter. Additive white Gaussian noise was included later during signal propagation. Figure 1b shows example waveforms and spectrograms for both call types, with and without Doppler-induced time warping.

#### Signal Propagation and Reception

To simulate signal reception at an array of microphones, I modelled a virtual sound source at position **s** = [*x*_*s*_, *y*_*s*_, *z*_*s*_] and evaluated its emission across the array.

The distance from the source to each microphone **m**_*i*_ was computed as:

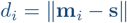

and the geometric propagation delay was:

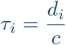

where *c* = 343 m*/*s is the speed of sound in air.

To realistically simulate the acoustic signal received at each microphone, I incorporated the following steps:

- **Source motion and Doppler resampling:** For a sound source moving at velocity **v**, the relative radial velocity *v*_*i*_ for each microphone was computed as:

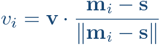

The corresponding Doppler factor *η*_*i*_ was used to time-warp the emitted waveform via rational resampling:

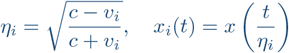

This compresses or stretches the signal depending on whether the source is approaching or receding, enabling the evaluation of Doppler-induced waveform distortion relevant to actively flying bats.

- **Spectral attenuation due to air absorption:** To account for distance-dependent frequency loss, the received signal was filtered in the frequency domain with an attenuation function:

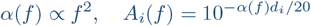

applied to the magnitude spectrum of the resampled signal.

- **Amplitude scaling for spherical spreading:** The resampled and attenuated signal was scaled by 1*/d*_*i*_ and normalised with respect to the first microphone.
- **Motion-induced delay correction:** A first-order correction term due to source motion was added to the geometric delay:

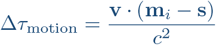

so that the total effective delay becomes *τ*_*i*_ = *d*_*i*_*/c* + Δ*τ*_motion_.

- **Fractional delay filtering:** The total delay was applied to each signal using a highresolution fractional delay filter implemented in the time domain.

These steps yielded a multichannel signal matrix with realistic propagation effects—including Doppler compression/stretching, distance-dependent spectral loss, amplitude attenuation, and sub-sample delay alignment—providing a controlled simulation framework for studying sound reception under dynamic acoustic conditions.

Note that all simulations were performed with *v* = 0 to evaluate localisation accuracy under stationary source conditions. The impact of source motion on localisation accuracy is discussed separately. The simulation tool includes methods to model motion-induced Doppler and delay effects for dynamic scenarios.

### 2.2 Localisation Methods

#### Time Difference of Arrival (TDoA) Estimation

TDoAs were extracted by computing the cross-correlation between the received signal at each microphone and a reference microphone (typically **m**_1_). The time delay Δ*τ*_*i*1_ was obtained as the lag that maximised the cross-correlation:

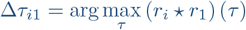

where *r*_*i*_ is the received signal at microphone *i*. Subsample interpolation was applied to refine delay estimates.

#### Multilateration

Given the TDoAs and known microphone positions, source localisation was formulated as a nonlinear least squares problem. The estimated source position ŝ was found by minimising the error between the observed and predicted TDoAs:

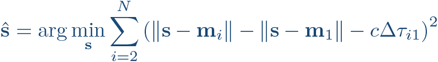

This was solved using MATLAB’s lsqnonlin optimiser, with **m**_1_ as the reference microphone.

#### Performance Metrics

For each simulation run, the localisation accuracy was assessed using:

- **Positional error**: Euclidean distance between true and estimated source positions,

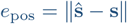

- **Angular error**: Difference in azimuth and elevation between ŝ*−***m**_1_ and **s***−***m**_1_, expressed in degrees.

### 2.3 Grid Sweep and Data Collection

To evaluate spatial accuracy, the simulation was run over a 3D grid spanning *x, y, z ∈* [*−*5, 5] m in 0.5 m increments, yielding a dense sampling of 3D space. At each point:

1. A virtual bat call was emitted.
2. Signals were propagated and received.
3. The source was localised via TDoA and multilateration.

For each array configuration, results including true/estimated coordinates, TDoAs, and error metrics were saved in structured CSV files for downstream analysis.

### 2.4 Simulation of Localisation Error due to Source Motion

To evaluate the effect of source velocity on localisation accuracy, I simulated bat calls emitted from a fixed source position while varying the source velocity. Two types of calls were tested: a broadband frequency-modulated (FM) chirp and a narrowband constant-frequency (CF) tone. Both call types were generated with a duration of 50 ms and an SNR of 60 dB, and were propagated to a tetrahedral four-microphone array (edge length: 0.5 m) positioned in space. The call signal was shaped using a Hanning window with 50% padding, and Doppler-induced time warping was introduced by resampling based on the instantaneous radial velocity of the source along the x-axis. The velocity was varied from *−*10 to +10 m/s in 0.1 m/s steps, simulating both approaching and receding trajectories.

A sampling rate of 192 kHz was used, and each call was localised using a time-differenceof-arrival (TDoA) approach based on cross-correlation. Ground-truth azimuth and elevation were computed from the vector connecting the reference microphone to the source. Angular and positional localisation errors were computed for each velocity condition. Simulations were repeated for both FM and CF calls to quantify their respective sensitivity to velocity-induced Doppler effects, which also effectively introduced phase distortions due to relative motion. A fixed random seed was used to ensure consistent noise generation across runs. The resulting errors were plotted to visualise the relationship between source velocity and localisation accuracy for each call type.

### 2.5 Statistical Analysis

To compare the performance of array configurations, the positional and angular errors were first partitioned into:

- **Inliers**: Positional errors *<* 10 cm *and* angular deviation *<* 2^*°*^
- **Outliers**: all others

For each group and configuration, summary statistics (mean, median, standard deviation) were computed.

A two-way analysis of variance (ANOVA) was performed with factors:

- *Configuration* (Tetrahedral, Square, Pyramid, Octahedral)
- *Error class* (Inlier, Outlier)

Post-hoc comparisons using Tukey’s HSD test identified statistically significant differences between pairs of array types. Visual summaries (boxplots, histograms, spatial error heatmaps) were generated to highlight error distributions and localisation bias patterns.

## 3 RESULTS

### 3.1 Microphone Array Configurations and Simulation Overview

Figure 1 illustrates the three microphone array geometries employed: the tetrahedron, planar square, and pyramid configurations. Each configuration consists of four microphones arranged to explore different spatial sampling strategies for three-dimensional localisation of ultrasonic signals. The synthesised bat call waveform and spectrogram (Figure 1b) confirm the generation of frequency-modulated signals characteristic of echolocating bats. An example localisation instance (Figure 1c) visually demonstrates the array layout, true source position, and estimated source location based on the time difference of arrival (TDoA) method, providing a baseline understanding of the simulation environment.

### 3.2 Spatial Distributions of Localisation Errors

Figures 2 and 3 present three-dimensional scatter plots of positional and angular localisation errors, respectively, for all tested microphone arrays. Positional errors are measured as the Euclidean distance between the true and estimated source positions, while angular errors represent discrepancies in azimuth and elevation angles from the array reference microphone.

**Figure 2:**
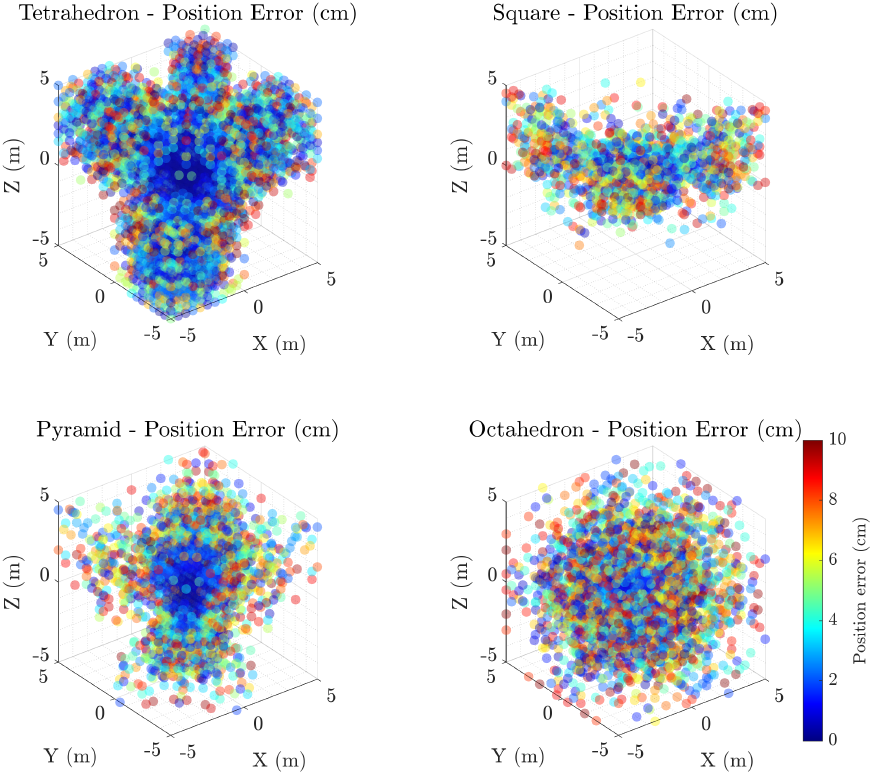
Spatial distributions of position localisation errors (in centimeters) for four different microphone array configurations used in this study: (a) Tetrahedron, (b) Square (Planar), (c) Pyramid, and (d) Octahedron. Each subplot presents a 3D scatter plot of the localisation error magnitude at various source positions within the measurement volume. Errors are color-coded with a colorbar indicating the scale, capped at 10 cm to highlight regions of high accuracy. The Tetrahedron and Pyramid configurations exhibit more concentrated clusters of low-error points near the origin, indicating higher localisation precision in the near field, whereas the Square (Planar) and Octahedron configurations display more dispersed error distributions. The Octahedron, with its larger spatial extent, shows increased variability and generally higher errors, suggesting trade-offs in array design for 3D localisation tasks.

**Figure 3:**
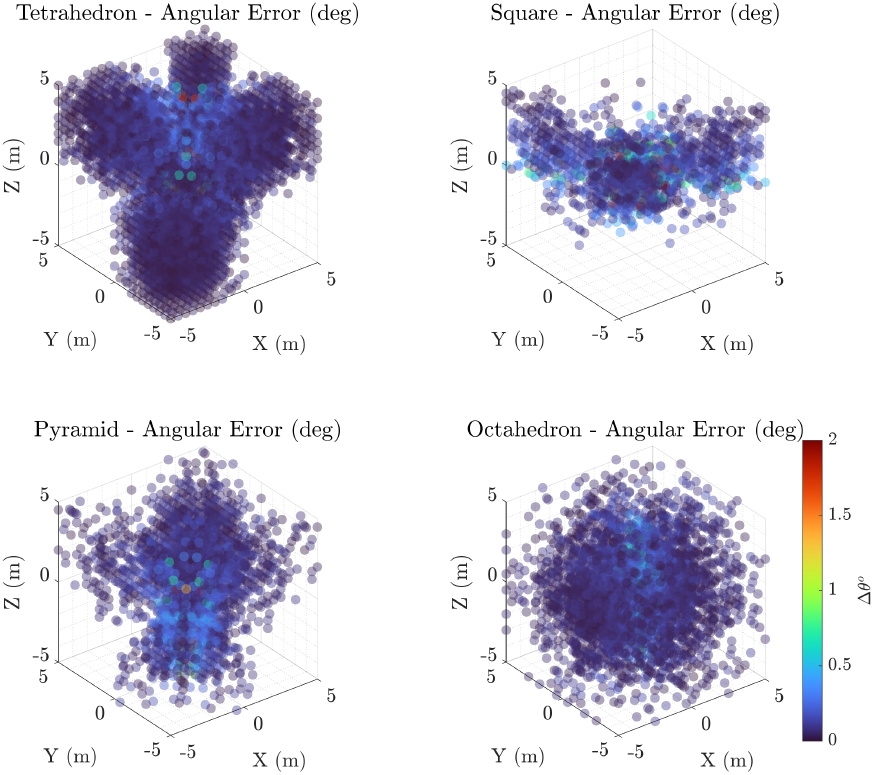
Spatial distributions of angular localisation errors (in degrees) for four distinct microphone array geometries: (a) Tetrahedron, (b) Square (Planar), (c) Pyramid, and (d) Octahedron. Each subplot presents a 3D scatter plot illustrating the magnitude of angular error at various source positions throughout the measurement space. The errors are color-coded with a colorbar indicating angular error magnitude, clipped at 2^*o*^ to emphasize regions of high angular precision. The Tetrahedron and Pyramid configurations demonstrate concentrated areas of low angular error near the array origin, indicating robust directional estimation in close proximity, while the Square (Planar) configuration shows increased angular variability, particularly in certain spatial zones. The Octahedron configuration reveals more uniform and generally lower angular errors, reflecting the advantage of 3D angular localisation over a larger spatial domain with increased number of receivers.

Visually, the tetrahedral configuration exhibits the most compact error clusters, suggesting superior localisation precision compared to the planar square and pyramid arrays, which display broader error spreads. Angular error scatter plots reinforce this observation, with the tetrahedron showing consistently lower angular deviations, indicative of better directional accuracy. These spatial error patterns underscore the impact of array geometry on localisation performance.

### 3.3 Localisation Error Distributions

Histograms of localisation errors (Figure 4) quantitatively summarise the error distributions for each microphone configuration. The position error histograms reveal that the tetrahedral array attains the lowest average positional error, with a sharper peak near zero error, while planar arrays demonstrate longer tails, indicating higher error variability. Angular error histograms mirror this trend; the tetrahedron consistently yields smaller angular deviations, whereas the pyramid and planar square exhibit wider angular error spreads. These histograms confirm the visual trends from the scatter plots and provide statistical context for the error magnitudes observed.

**Figure 4:**
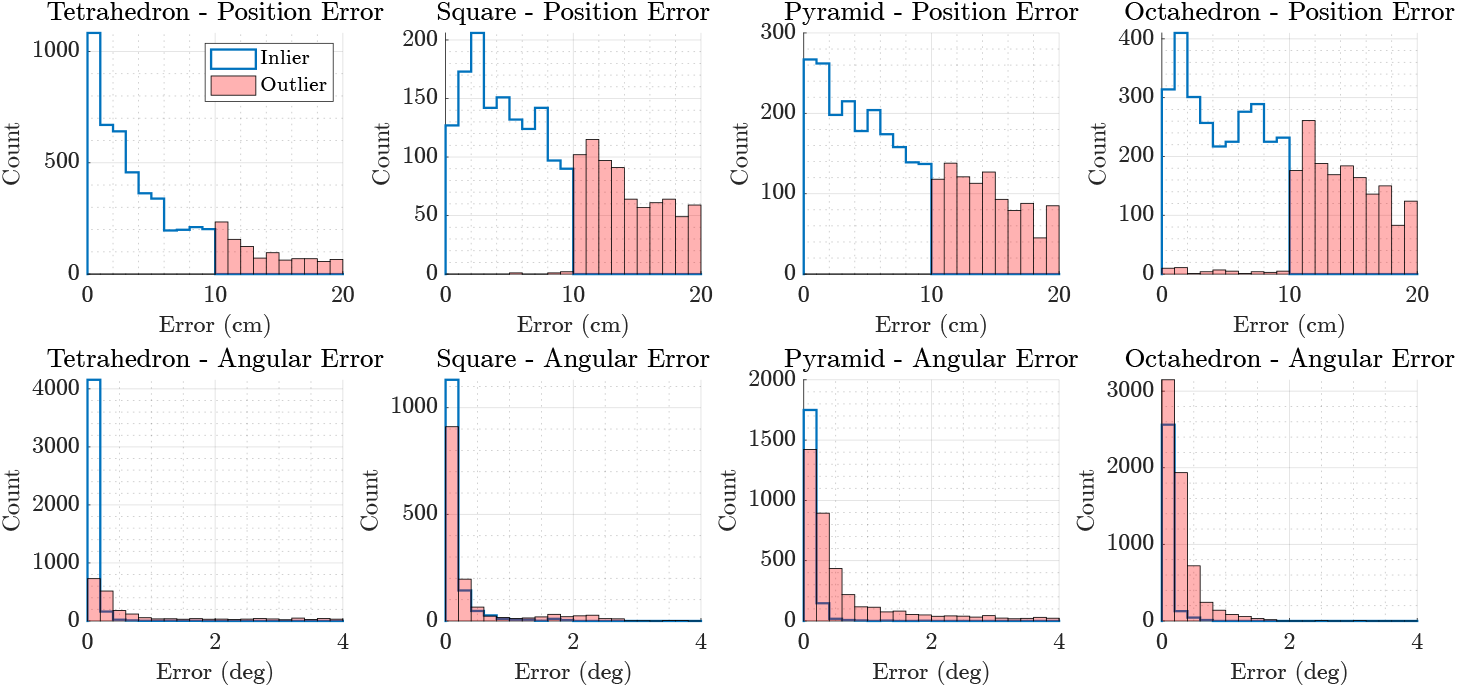
Histograms of localisation error distributions for four microphone array configurations: Tetrahedron, Square (Planar), Pyramid, and Octahedron. The top row displays position error (in centimeters), while the bottom row shows angular error (in degrees). Inliers are erros < 10 cm *and* 2^*o*^. Each histogram differentiates between *inliers* (blue lines/bars), representing errors below the defined thresholds, and *outliers* (red shaded bars), indicating localisation failures or larger errors beyond threshold limits. Position errors exhibit a strong concentration near zero for Tetrahedron and Pyramid arrays, illustrating higher spatial accuracy, whereas Square and Octahedron arrays show broader and more frequent large errors. Angular error distributions follow a similar trend, with Tetrahedron and Pyramid arrays achieving superior directional precision, with the highest angular precision achieved by the Octahedron. This distinction between inliers and outliers highlights the robustness and reliability of each microphone geometry for precise localisation tasks.

### 3.4 Localisation Error Summary by Inliers and Outliers

Boxplots in Figure 5 further delineate localisation accuracy by categorising results into inliers (errors within 10 cm and 2 degrees) and outliers. The inlier boxplots show that the tetrahedral array achieves the lowest median and spread for both positional and angular errors, reinforcing its higher precision. Conversely, outlier distributions highlight the presence of large errors predominantly in the planar square and pyramid arrays, demonstrating that these configurations are more prone to significant localisation failures.

**Figure 5:**
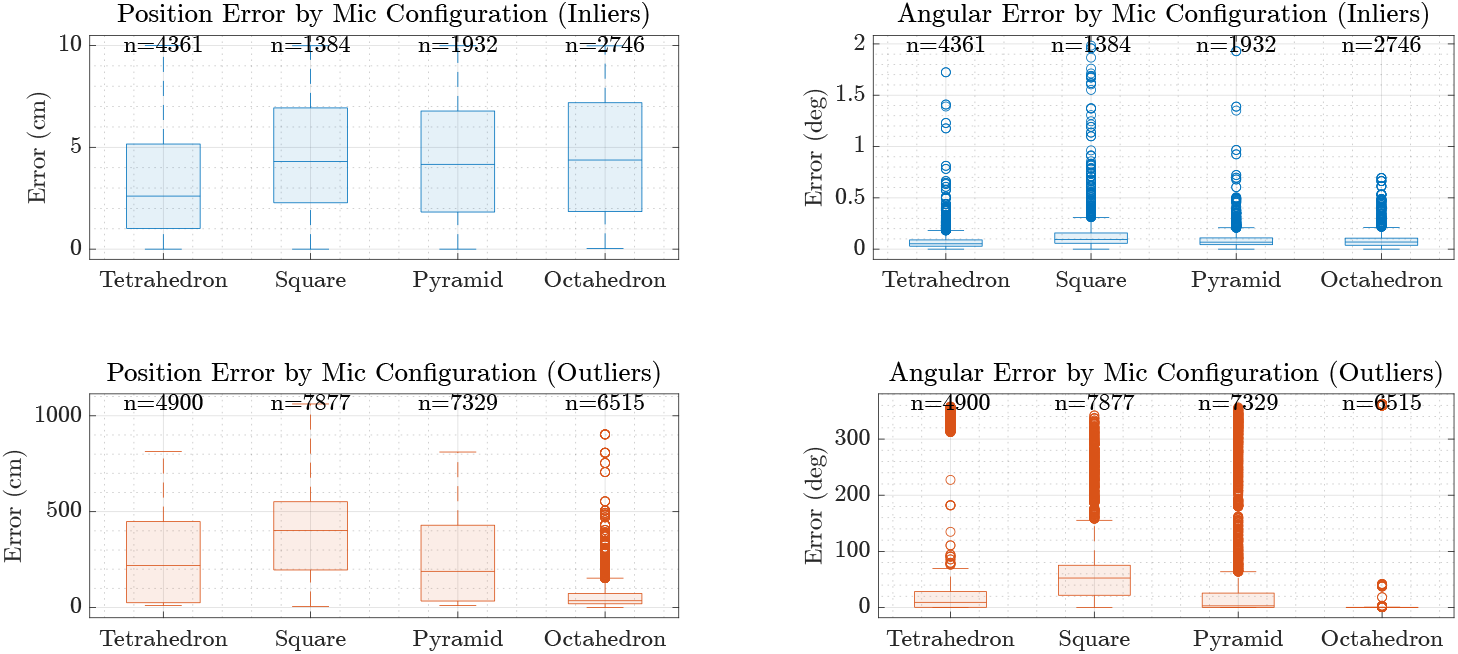
Boxplots illustrating the distributions of localisation errors separated into *inliers* and *outliers* for four different microphone array configurations: Tetrahedron, Square (Planar), Pyramid, and Octahedron. The top row presents the positional (centimeter) and angular (degree) errors for inliers, while the bottom row shows the corresponding errors for outliers. The Tetrahedron configuration demonstrates a superior balance, achieving consistently lower median and mean positional and angular errors among the inliers, indicating robust and reliable localisation performance across both spatial and directional domains. In contrast, the Octahedron array, while exhibiting comparable positional error distributions, excels in angular precision with notably lower angular errors and fewer extreme outliers in angular localisation. The Square (Planar) and Pyramid configurations show comparatively larger spreads and higher median errors, particularly in the positional domain for outliers, suggesting limitations in resolving precise locations in complex acoustic environments. These differences highlight the trade-offs between array geometries, where the Tetrahedron offers a well-rounded compromise, and the Octahedron provides specialized angular accuracy, relevant for applications prioritizing directional precision.

Sample size annotations above each box provide transparency on the number of valid measurements, illustrating that while all arrays capture substantial valid localisations, the quality varies significantly across configurations.

### 3.5 Positional Error Contour Maps Across Elevation Slices

Figure 6 presents contour maps of positional localisation errors across spatial grids segmented by elevation slices (Z-values). These heatmaps, clipped at 50 cm for clarity, visually communicate spatial error patterns within the testing volume. The tetrahedral configuration consistently shows concentrated low-error regions across most slices, while planar square and pyramid arrays exhibit larger areas with elevated errors, especially at negative elevations.

**Figure 6:**
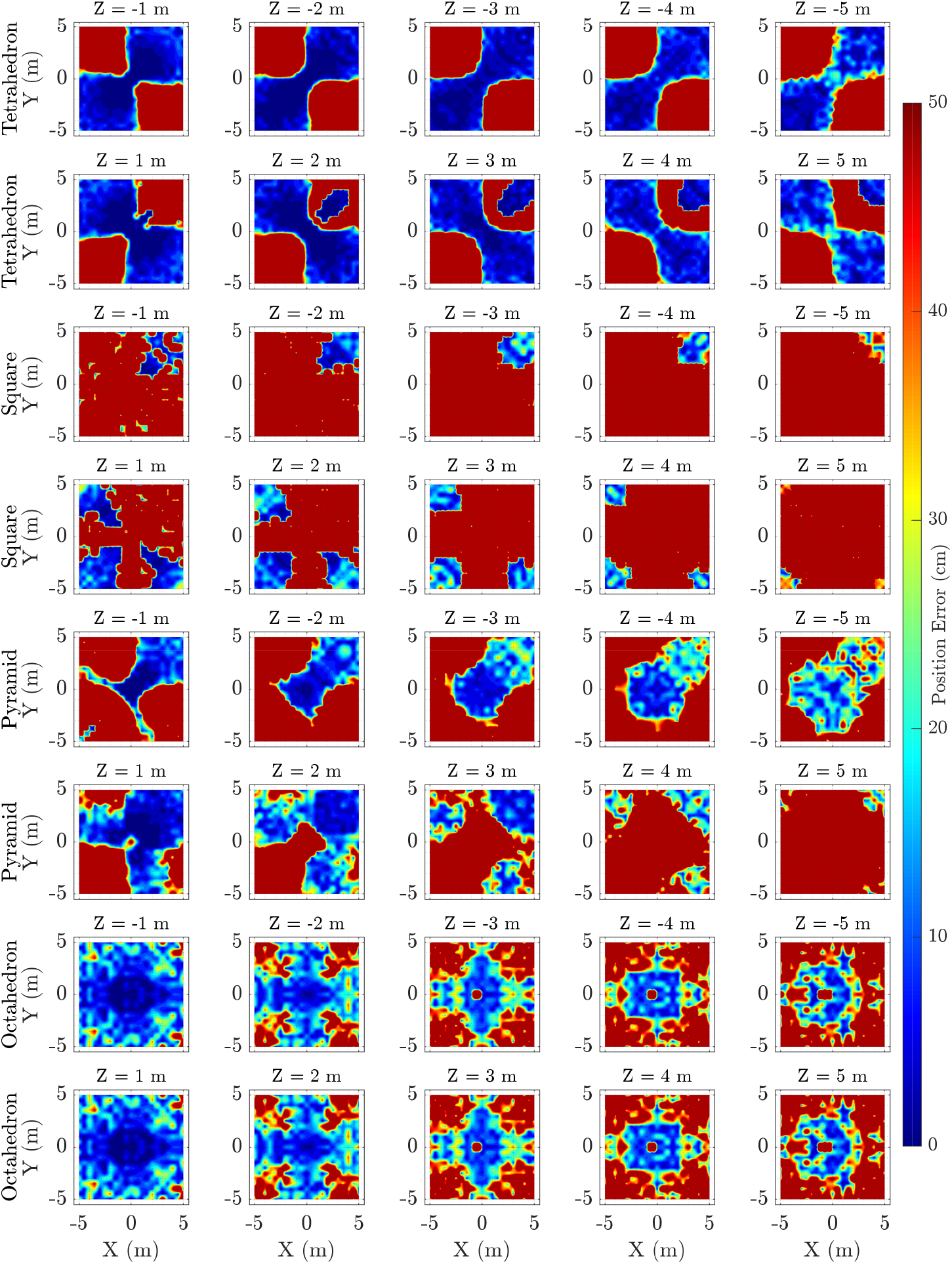
Heatmaps of positional localisation error (cm) for different microphone array configurations across multiple elevation slices (*Z* = *±* 1 to *±* 5 m). Each row represents a microphone configuration: Tetrahedron, Square (Planar), Pyramid, and Octahedron (top to bottom). The columns correspond to discrete elevation planes. The Tetrahedron and Pyramid arrays show relatively compact regions of low error near the center, with errors increasing towards the edges. The Square (Planar) configuration demonstrates notably larger regions of high positional error, especially at higher elevation slices, indicating reduced 3D localisation capability. The Octahedron, while more spatially uniform, exhibits increased errors at the edges, possibly reflecting the influence of array geometry on far-field precision. Overall, these heatmaps reveal how array geometry and source elevation influence localisation accuracy, guiding optimal design considerations for 3D microphone arrays.

The paired-row layout for negative and positive elevations highlights how errors vary with vertical position, revealing potential array blind spots or coverage limitations inherent in the planar geometries. These spatial visualisations offer critical insight into where and why each configuration excels or struggles.

### 3.6 Angular Error Contour Maps Across Elevation Slices

Similar contour maps of angular localisation errors are depicted in Figure 7, with error clipping at 10 degrees for enhanced interpretability. The tetrahedral array maintains low angular error regions broadly, indicating robust directional estimation throughout the volume. The planar square and pyramid configurations again display wider zones of elevated angular error, reflecting the geometric constraints and reduced 3D coverage of these arrays.

**Figure 7:**
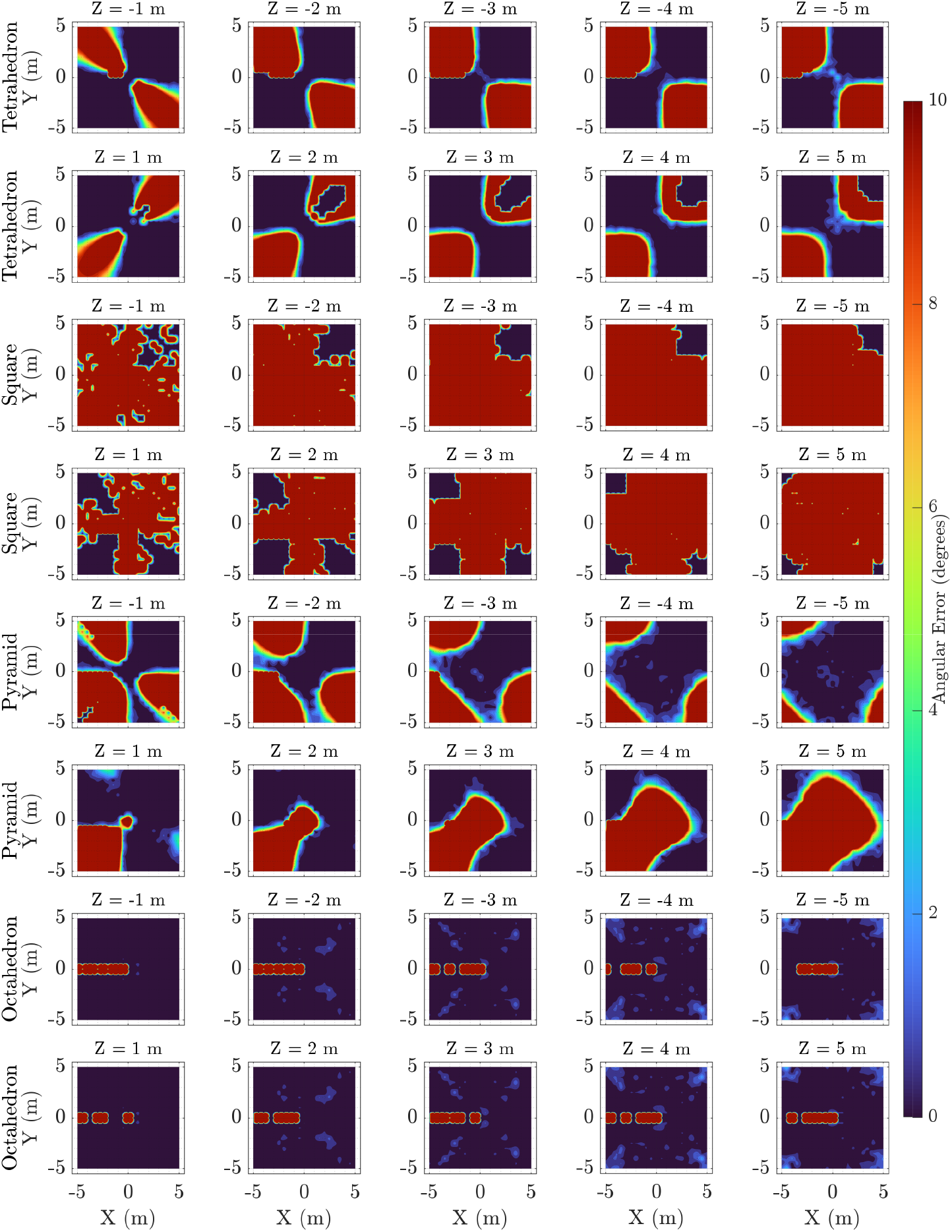
Contour maps of angular localisation error (degrees) for different microphone array configurations across a series of elevation slices. The Tetrahedron and Pyramid configurations exhibit localized regions of low angular error, particularly near the array center, with errors increasing towards the edges and at higher elevations. The Square (Planar) configuration demonstrates significantly larger angular errors and less spatial uniformity, indicating limitations in vertical spatial resolution. The Octahedron achieves consistently low angular errors across all slices, reflecting its superior ability for angular localisation precision. Overall, these spatial error maps highlight how microphone array geometry and elevation influence the accuracy of angular localisation, providing valuable insights for optimizing array designs tailored to specific bioacoustic monitoring scenarios.

By analysing these angular error distributions, it becomes evident that three-dimensional microphone placements that avoid coplanarity, such as the tetrahedron, provide superior angular resolution for ultrasonic source localisation.

## 3.7 Statistical Analysis of Localisation Performance

Table 1 quantitatively summarises localisation errors by microphone configuration and grouping into inliers and outliers. The tetrahedral array demonstrates significantly lower mean and median positional and angular errors among inliers, consistent with visual data. In contrast, the planar square and pyramid arrays show higher error statistics and greater variability.

**Table 1:** Summary statistics of positional and angular localisation errors (in centimeters and degrees, respectively) for different microphone array configurations. Results are split by inlier and outlier groups based on threshold criteria. The tetrahedron configuration achieves the best overall positional accuracy among inliers, while the octahedron offers superior angular precision. Outliers demonstrate considerably larger errors, highlighting the importance of filtering for accurate localisation. Notably, the values did not change for simulation runs with different call types and call durations.

| Config | Group | Pos. Mean | Pos. Med | Pos. Std | Ang. Mean | Ang. Med | Ang. Std |
| --- | --- | --- | --- | --- | --- | --- | --- |
| Octahedron | Inlier | 4.68 | 4.37 | 2.93 | 0.09 | 0.07 | 0.09 |
| Octahedron | Outlier | 63.30 | 35.57 | 84.40 | 6.51 | 0.21 | 46.61 |
| Pyramid | Inlier | 4.40 | 4.16 | 2.83 | 0.10 | 0.07 | 0.12 |
| Pyramid | Outlier | 242.12 | 188.64 | 213.33 | 35.75 | 3.05 | 79.55 |
| Square | Inlier | 4.54 | 4.30 | 2.74 | 0.17 | 0.09 | 0.24 |
| Square | Outlier | 376.98 | 401.81 | 241.20 | 60.76 | 52.52 | 60.52 |
| Tetrahedron | Inlier | 3.34 | 2.61 | 2.77 | 0.07 | 0.05 | 0.09 |
| Tetrahedron | Outlier | 254.91 | 218.95 | 221.15 | 35.06 | 9.28 | 81.38 |

ANOVA results indicate statistically significant differences between microphone configurations and error groups, with post-hoc tests confirming the tetrahedral geometry’s superior performance is statistically robust. These analyses provide rigorous validation for selecting array geometries optimised for three-dimensional ultrasonic localisation.

The comprehensive simulation and statistical analyses establish the tetrahedral microphone array as the optimal configuration for precise three-dimensional localisation of high-frequency ultrasonic signals. Its spatial arrangement yields improved positional and angular accuracy across the evaluation volume, outperforming planar and pyramidal alternatives. These findings guide future array design for bioacoustic monitoring and related applications requiring precise ultrasonic source localisation.

### 3.8 Effect of Call Duration and Type

To evaluate the influence of call characteristics on localisation accuracy, I compared performance across simulated frequency-modulated (FM) and constant-frequency (CF) calls, at 5, 10 and 50 ms signal durations. The analysis revealed no systematic differences in localisation accuracy between call types under these controlled conditions. Despite the spectral differences between FM and CF signals, localisation errors remained comparable across both types, indicating that the acoustic structure of the call alone does not significantly affect TDoA-based localisation performance.

### 3.9 Effect of Source Velocity on Localisation Accuracy

The analysis demonstrated that localisation performance is strongly affected by source velocity, with both positional and angular errors increasing markedly as velocity magnitude increases. As shown in Figure 8, this degradation was broadly symmetric for positive and negative velocities but was particularly severe for constant-frequency (CF) calls at moderate speeds.

**Figure 8:**
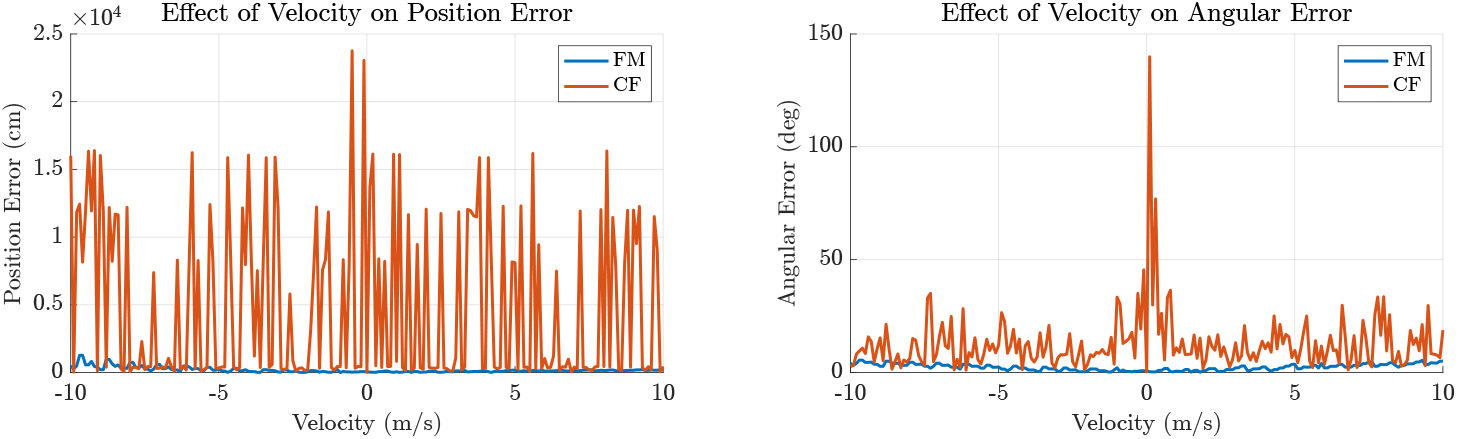
Effect of source velocity on localisation error for FM and CF calls. Simulations using a tetrahedral microphone array demonstrate that localisation accuracy degrades with increasing source velocity due to Dopplerinduced distortion of the received signal. Frequency-modulated (FM) calls (10 ms, 25–80 kHz sweep) remain robust across the tested velocity range, exhibiting minimal position and angular error. In contrast, constant-frequency (CF) calls (100 ms, 80 kHz tone), representative of Hipposiderid and Rhinolophid bats, show highly unstable localisation at moderate velocities, with large spikes in both position and angular error. This reflects their higher susceptibility to Doppler time-warping and phase distortion over long durations. These results highlight the importance of call structure in motion-compensated acoustic localisation and demonstrate that FM calls, like those used by many Vespertilionid bats, offer significantly greater resilience to velocity-induced errors.

FM calls (10 ms duration, 25–80 kHz sweep), representative of Vespertilionid bats, remained consistently robust across the full range of tested velocities, showing only modest increases in localisation error. In contrast, CF calls (100 ms duration, 80 kHz tone), exhibited substantial spikes in both position and angular error. This suggests that the Doppler-induced time warping and phase distortions during long-duration tonal calls are not well compensated by the TDoA-based localisation algorithm, leading to unstable and unreliable estimates.

These results indicate that call type has a substantial influence on localisation performance under motion, with FM sweeps offering greater resilience to velocity-induced distortion. Consequently, acoustic localisation systems intended for tracking CF-emitting species may require additional correction strategies to achieve reliable performance under natural movement conditions.

## 4 DISCUSSION

Even though a number of studies have employed microphone arrays with varying numbers of elements – typically arranged in two-dimensional planar configurations– for studying bat echolocation behaviour, including field-based recordings of *Myotis daubentonii* by Surlykke et al. [21], beam shape analysis of *Carollia perspicillata* by Brinkløv et al. [22], spectral effects of environmental conditions by de Frémond et al. [23], and flight speed analyses by Jakobsen et al. [24], it is noteworthy that most of these studies either do not report localisation accuracy, or rely on manual filtering of detections to retain only positions that conform to expected trajectories. The absence of a standardised benchmark for evaluating localisation precision across array designs and signal types remains a significant gap in the field, limiting the reproducibility and comparability of results obtained via TDoA-based localisation methods.

This study explores the spatial distribution of three-dimensional positional and angular localisation accuracies for three distinct microphone array configurations, each comprising four microphones, and one with six in an octahedral form. The developed simulation framework, encompassing signal generation, reception, and localisation, is a crucial tool for characterising and designing 3D microphone arrays optimised for ultrasonic source localisation. By evaluating localisation precision using the implemented time difference of arrival (TDoA) method, detailed error matrices are generated, which provide insight into expected real-world performance and may be used in localisation analyses of field data.

In evaluating the practical implications of localisation accuracy, it is instructive to contextualise the positional and angular errors relative to typical bat flight speeds and echolocation call characteristics. Considering a conservative error threshold of 10 cm for positional accuracy, I examine its significance at nominal bat flight velocities of 2, 3, 4, and 5 m/s. The time *t* required for a bat to traverse this 10 cm spatial resolution can be calculated by the simple relation:

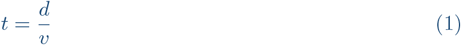

where *d* = 0.1 m is the error limit distance and *v* is the velocity. Substituting values yields: 50 ms, ≈ 33 ms, 25 ms, 20 ms, respectively.

These time intervals correspond to call rates of approximately 20, 30, 40, and 50 Hz, respectively, assuming calls are spaced to match the bat’s progression through the error zone. This alignment suggests that the positional resolution of 10 cm fits well within the natural echolocation call timing of many species, enabling effective tracking without significant aliasing or temporal ambiguity.

For angular localisation, an error of 1 degree corresponds to a linear spatial uncertainty that depends on the distance *r* from the microphone array. This can be approximated by the arc length formula:

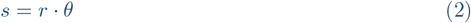

where *θ* is the angular error in radians. Converting 1^*°*^ to radians (*θ* = *π/*180) and calculating for distances *r* = 1, 2, 3, 4, 5 m gives:

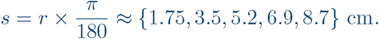

Thus, a 1-degree angular error results in linear errors under 9 cm at typical localisation distances, which is within or close to the observed positional error bounds.

Furthermore, typical bat echolocation calls have durations on the order of 5 ms. The linear distance a bat covers during one call is:

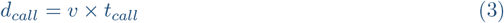

For the aforementioned velocities and a call duration *t*_*call*_ = 5 ms, it corresponds to 1, 1.5, 2, 2.5 cm, respectively.

These distances are significantly smaller than the positional error threshold, implying that localisation errors on the order of 10 cm will not obscure the detection of fine-scale behavioural changes during individual calls but rather affect spatial resolution across multiple calls.

Given the mean angular errors obtained, especially for the tetrahedral configuration, where angular precision often remains below 1 degree for inliers, the TDoA localisation method demonstrates high reliability within biologically relevant spatial and temporal scales. This precision is crucial for detailed behavioural studies where accurate tracking of echolocating bats’ positions and orientations directly informs ecological and neuroethological interpretations.

Overall, these considerations highlight that the defined positional and angular error limits are well aligned with bat locomotion and echolocation call dynamics, affirming the practical utility of the developed localisation system for in-field bioacoustic studies.

The echolocation calls of bats, despite the rapid attenuation often observed at the start frequencies of typical frequency-modulated (FM) calls, present distinct advantages for acoustic localisation compared to many bird vocalisations. These advantages arise from several inherent characteristics of bat calls and the acoustic environment in which they occur. Bat calls are broadband signals, generally very short in duration, ranging from approximately 0.5 to 10 milliseconds for FM bats [8], and up to 100 milliseconds for constant-frequency (CF) bats [25]. This brevity reduces the likelihood of temporal overlap, even when multiple bats are flying and vocalising in close proximity.

The probability of overlapping bat calls in the time domain is exceedingly small, as it requires a highly precise coincidence: bats must vocalise simultaneously, and their emitted calls must arrive at the microphones with matching time-of-arrival differences. Such precise temporal and spatial alignment implies that the bats are receiving near-simultaneous stimuli and reacting synchronously, a condition that is statistically rare in natural flight behaviour. Consequently, time-domain resolution of bat calls using cross-correlation techniques is highly reliable.

Furthermore, the ultrasound frequency band used by bats is typically free from significant environmental noise, except for some insect species such as cicadas and katydids during specific seasons [26]. This low ambient noise floor enhances the signal-to-noise ratio (SNR) for bat calls, enabling the detection of distinct cross-correlation peaks in the received signals. These peaks can often be resolved with microsecond precision, facilitating highly accurate TDoA measurements essential for localisation.

Together, these factors make bat echolocation calls particularly well suited for robust and precise acoustic localisation using microphone arrays, as the combined temporal sparsity, broadband nature, and quiet ultrasonic environment simplify signal separation and reduce ambiguities commonly encountered in the localisation of other animal vocalisations.

Figure 1a presents the four microphone array geometries tested: the tetrahedron, planar square, pyramid, and octahedron. These configurations differ in spatial microphone distribution: the tetrahedron and octahedron are fully three-dimensional and symmetrical, while the square array is strictly planar, and the pyramid offers a compromise with limited vertical spread. These geometric differences are expected to influence each array’s capacity to resolve three-dimensional source positions, particularly along the vertical axis. The simulated bat call waveform and spectrogram (Figure 1b) represent the frequency-modulated ultrasonic signals used in all tests, while Figure 1c illustrates the typical localisation setup.

The 3D scatter plots of positional and angular errors (Figures 2 and 3) reveal distinct localisation behaviours among the arrays. The tetrahedral array consistently clusters positional and angular errors near zero, indicating superior spatial resolving power across all axes. The octahedron also achieves relatively compact error clusters, particularly in angular estimation, highlighting the advantage of its high spatial symmetry and increased microphone count. In contrast, both the square and pyramid configurations produce more dispersed error distributions, particularly in the elevation (z-axis), consistent with their more limited vertical sensor separation. This dispersion reflects reduced triangulation precision in directions not well sampled by the microphone geometry.

Across all configurations, angular errors tend to be lower than positional errors, suggesting that direction estimation (based on TDoA differences) is inherently more robust than precise range estimation. Notably, the tetrahedron and octahedron show tighter angular error clusters compared to the planar and pyramidal arrays, reinforcing the importance of 3D spatial coverage for accurate direction-of-arrival (DoA) estimation.

Histograms of localisation error (Figure 4) offer further quantitative insight. The tetrahedral and octahedral arrays exhibit sharp, narrow peaks near zero for both positional and angular errors, indicative of consistent performance. The octahedron, while marginally outperformed by the tetrahedron in overall positional accuracy, shows a notably tighter angular error histogram, suggesting it is particularly well-suited for applications prioritising directional tracking. By contrast, the square and pyramid arrays display broader error distributions and heavier tails, reflecting greater variability and more frequent large errors. These trends confirm that array dimensionality and symmetry play key roles in localisation reliability.

Figure 5 explores these differences in greater detail by distinguishing inliers—localisations with errors under 10 cm and 2^*°*^—from outliers. The tetrahedral array shows the lowest median and interquartile range for both error types among inliers, underscoring its robustness. Interestingly, the octahedron exhibits the lowest median angular error of all four configurations, despite slightly higher positional spread compared to the tetrahedron. This trade-off highlights its directional stability and potential advantage in applications where bearing estimation is prioritised over range. In contrast, the square and pyramid arrays show wider error distributions, especially among outliers, with the planar configuration performing poorest under spatially challenging conditions.

Elevation slice contour maps (Figure 6) provide a spatially resolved view of positional accuracy across the localisation volume. The tetrahedron maintains uniformly low error regions throughout the field, confirming its consistent coverage. The octahedron similarly shows broad low-error zones, albeit with slightly more variability across slices, likely due to geometric redundancy and overdetermined TDoA estimates. The square and pyramid arrays, by contrast, reveal distinct zones of poor accuracy—particularly at negative elevations—demonstrating blind spots resulting from limited vertical microphone distribution.

Angular error contour maps (Figure 7) support these conclusions. The tetrahedral and octahedral configurations both maintain low angular errors across most slices, while the square and pyramid geometries suffer from degraded performance at off-centre elevations. This again highlights the limitations of 2D or quasi-3D microphone layouts for full 3D localisation tasks.

The statistical summary in Table 1 affirms these visual analyses. Both the tetrahedral and octahedral arrays show significantly lower mean and median positional and angular errors than the square and pyramid configurations. Standard deviations are also smaller for these fully 3D arrays, indicating less variability. ANOVA tests confirm the statistical significance of these differences, with post-hoc comparisons ranking the tetrahedral array as the most balanced and consistently accurate configuration overall, and the octahedron as a strong alternative for angular estimation tasks.

The observed asymmetry in localisation accuracy—especially the decline at negative elevations—is attributable to the arrays’ orientation and geometric limitations. All arrays were modelled with their bases horizontal, a common field deployment choice. However, the directional nature of bat echolocation calls, combined with wavefront curvature and propagation direction, means that arrays lacking vertical microphone distribution are intrinsically less sensitive to sources located below the array plane. This insight emphasises the importance of deliberate array orientation in field settings and suggests that augmenting planar arrays with additional vertical elements could significantly improve spatial performance.

### 4.1 Effect of Source Motion During Call Emission

An important but often overlooked source of localisation error in TDoA-based systems arises when the sound source is in motion during the emission of a call. Echolocating bats emit calls while flying, and even small displacements during a single call can distort the signal’s propagation path to each microphone. This dynamic motion causes unequal phase shifts and time warping across channels that cannot be fully corrected using standard delay-based alignment, particularly when employing cross-correlation for TDoA estimation.

These effects are especially pronounced in bats that emit long-duration calls. Constant-frequency (CF) bats, for instance, often produce calls that are 20–100 ms in duration. While these signals are narrowband and spectrally simple, their extended duration makes them particularly susceptible to localisation inaccuracies. During the call, the bat’s position and velocity may both change, introducing a non-uniform phase distortion across microphones. This motion results in a complex interaction between geometric delay, Doppler shift, and potential spectrotemporal manipulations due to signal-reflections overlap, which compromises the precision of delay estimation and ultimately localisation accuracy [21, 27].

In any microphone array configuration, each transducer experiences a different relative velocity per dimension and angular orientation with respect to the moving bat. Consequently, the emitted call is warped differently at each microphone, violating the assumption that signals differ only by time delay. This introduces further errors in the cross-correlation peak and the resulting TDoA estimate. These sources of error may be how some studies involving localisation of Hipposiderid bats have reported worse performance using acoustic methods compared stereo-camera based appraoch [27–29].

In contrast, frequency-modulated (FM) calls are typically shorter in duration (1–10 ms), which reduces the window over which motion-induced distortion can occur. However, FM calls are more broadband and sensitive to phase coherence, making them vulnerable tol inconsistencies across channels caused by motion or angular displacement.

Figure 8 illustrates the impact of source motion during call emission on localisation accuracy, corroborating the mechanisms described above.

While this study primarily evaluated localisation accuracy under the assumption of a stationary source (*v* = 0), the simulation framework incorporates the ability to model time-varying Doppler and motion effects by resampling the emitted call based on the source’s relative velocity to each microphone. This allows investigation of motion-induced errors in both FM and CF contexts.

### 4.2 Localisation Accuracy Enhancement

The accuracy of acoustic localisation using time-difference-of-arrival (TDoA) fundamentally depends on array geometry, signal properties, and environmental conditions. While angular localisation is often precise, particularly in well-spaced microphone configurations, radial or distance accuracy is inherently less constrained in TDoA-based systems. This limitation arises because the difference in arrival times maps each pair of microphones to a hyperboloid surface of potential source locations. Without sufficient spatial diversity in microphone placement, particularly along the depth axis, these surfaces can intersect ambiguously, resulting in poor resolution in the radial direction [16, 30–32].

Highly symmetric or coplanar geometries, such as square or circular arrays, exacerbate this issue by producing geometrically degenerate TDoA patterns—where multiple source locations yield similar time differences—thereby increasing uncertainty and susceptibility to noise [33]. In contrast, asymmetric three-dimensional arrays, such as the tetrahedral configuration used in this study, break these symmetries and provide more unique TDoA signatures across space, enhancing overall localisation fidelity. The lack of coplanarity and increased volumetric spread of microphones help constrain the source location in all three dimensions, reducing ambiguity in both azimuth-elevation space and radial distance.

### 4.3 Limitations

Although this study provides a mathematically robust framework for evaluating array configurations prior to physical construction, and the accompanying software is feature-rich, extensible, and fully open-source, experimental validation of the predicted localisation accuracies remains necessary. Empirical tests may reveal additional factors that require further refinement or calibration of the models.

Importantly, this study adopts a heuristic approach, focusing not on exhaustively identifying the sources of localisation error, but rather on enabling task-specific array design through comparative evaluation. While identifying error sources is undoubtedly crucial for enhancing accuracy, doing so demands dedicated future investigations that extend beyond the scope of this work.

The simulations suggest a lower bound on localisation resolution in the range of 5–10 cm; however, this is inherently constrained by the chosen array scale, with all array arms fixed at 0.5 m in this study. Alternative geometries or larger arrays may shift these performance bounds and should be explored in future work.

Moreover, the current implementation assumes idealised recording conditions. Real-world scenarios introduce additional challenges such as environmental noise, multipath reflections, and variability across microphones. Field recordings, in particular, are vulnerable to signal distortions beyond the researcher’s control. In addition to motion-induced effects discussed earlier, reflections from the ground, vegetation, nearby objects, or even inadequately insulated equipment can modulate the signal. These effects are especially problematic for long-duration calls, which are more likely to overlap with delayed echoes. For example, a 10 ms call can be significantly affected by ground reflections if the microphone is positioned 1–2 m above the surface. These factors complicate the reliable localisation of wild bats and underscore the importance of accounting for environmental acoustic interactions in experimental design and data interpretation.

Future improvements may include adaptive or modular array architectures with reconfigurable geometry or self-calibrating capabilities to maintain robust localisation performance across changing field conditions. Furthermore, integrating probabilistic frameworks or machine learning-based inference could enhance the system’s ability to model uncertainty and operate effectively in acoustically complex environments.

In parallel, further work is in progress to develop cost-effective, field-deployable array systems using custom-built, microcontroller-based multichannel ultrasonic recorders that are highly portable and energy-efficient [34]. Such tools not only facilitate practical deployment in field experiments but also support the continued development and real-world validation of the methods described here.

## 5 CONCLUSIONS

This study presents a modular, open-source simulation framework for evaluating the localisation accuracy of 3D microphone arrays under diverse signal and motion conditions. Using the *Array WAH* toolkit, I systematically compared several standard geometries and demonstrated how spatial configuration influences both angular and positional accuracy. Tetrahedral and octahedral arrays showed the highest robustness across conditions, offering design guidance for compact yet effective field-deployable arrays.

My results highlight the influence of source motion during call emission, particularly in CF bats emitting long-duration calls. While FM and CF calls yielded comparable localisation errors under the simulated conditions, motion-induced distortions become increasingly relevant with higher velocities and asymmetric geometries. These findings stress the importance of choosing configurations that reduce ambiguity regions and enhance phase separability across microphones.

Despite the idealised assumptions, the framework provides a reproducible, extensible platform to inform real-world deployment. Limitations such as environmental noise, multipath effects, and device variability underscore the need for experimental validation. Ongoing work focuses on cost-effective field deployment using a custom microcontroller-based ultrasound recorder [34], enabling in situ testing of the localisation principles described here.

Overall, the toolkit provides a versatile method to optimise sensor geometry and supports custom design for bioacoustic and ecological applications.

## 6 DECLARATIONS

### Ethics approval and consent to participate

Not applicable. This study did not involve experiments on animals or humans.

### Consent for publication

Not applicable.

### Availability of data and materials

The source code for the Array WAH project, including MATLAB scripts for evaluating 3D localisation performance across microphone array geometries, is publicly available at: https://github.com/raviumadi/Array_WAH.git

Additionally, a data repository has been archived on Zenodo and is available at: https://doi.org/10.5281/zenodo.15691371

### Author contributions

**Ravi Umadi:** Conceptualisation, Methodology, Software, Formal analysis, Investigation, Visualisation, Validation, Writing – original draft, Writing – review & editing, Project administration, Resources.

## Acknowledgements

I would like to extend thanks to my colleagues at the Lehrstuhl für Zoologie, Weihenstephan for reviews and comments. I sincerely thank the reviewers for their constructive feedback, which has significantly contributed to refining the manuscript and expanding its scope and applicability. I would also like to thank Irina Tolkova for her valuable comments on an earlier draft of the manuscript.

